# Cell-derived hexameric β-amyloid: a novel insight into composition, self-assembly and nucleating properties

**DOI:** 10.1101/2020.12.15.422916

**Authors:** Devkee M. Vadukul, Céline Vrancx, Pierre Burguet, Sabrina Contino, Nuria Suelves, Louise C Serpell, Loïc Quinton, Pascal Kienlen-Campard

## Abstract

A key hallmark of Alzheimer’s disease (AD) is the extracellular deposition of amyloid plaques composed primarily of the amyloidogenic amyloid-β (Aβ) peptide. The Aβ peptide is a product of sequential cleavage of the Amyloid Precursor Protein (APP), the first step of which gives rise to a C-terminal Fragment (C99). Cleavage of C99 by γ-secretase activity releases Aβ of several lengths and the Aβ42 isoform in particular has been identified as being neurotoxic. The misfolding of Aβ leads to subsequent amyloid fibril formation by nucleated polymerisation. This requires an initial and critical nucleus for self-assembly. Here, we identify and characterise the composition and self-assembly properties of cell-derived hexameric Aβ42 and show its nucleating properties which are dependent on the Aβ monomer availability. Identification of nucleating assemblies that contribute to self-assembly in this way may serve as therapeutic targets to prevent the formation of toxic oligomers.

## Introduction

Alzheimer’s disease (AD) is a neurodegenerative disease characterised by the deposition of extracellular amyloid plaques in the brain which are primarily composed of the self-assembled amyloid-β (Aβ) peptide[1]. The self-assembly process of Aβ has been the focus of much research, however, it is still unclear how this relates to disease pathology. Aβ is a product of sequential cleavage of the Amyloid Precursor Protein (APP) in the amyloidogenic pathway where APP is first cleaved by β-secretase at the N-terminus of Aβ. The two products of this are soluble APP-β (sAPPβ) and a C-terminal fragment (CTF) consisting of 99 amino acids (C99). C99 is further cleaved by γ-secretase beginning with a proteolytic cut at the ε site to free the C-terminal APP intracellular domain (AICD) and release Aβ of varying lengths ranging from Aβ49-38[2-4].

Although extracellular Aβ plaques found in AD brains are primarily composed of highly ordered cross-β mature amyloid fibrils[5-7], soluble forms of Aβ have shown a much greater correlation with cognitive decline and neurodegeneration in AD patients[8,9]. Due to this, it is now widely accepted that pre-fibrillar spherical/globular Aβ oligomers are neurotoxic entities. In particular, several Aβ42 oligomers of different assembly sizes and conformations have been identified as being cytotoxic[7,10-18].

Oligomers are formed as intermediary assemblies during amyloid formation, the mechanism of which is via nucleated polymerisation[19]. There is first the nucleation or lag phase where the monomer precursor is either in an unfolded, partially folded or natively folded state and undergoes usually unfavourable self-association to form nuclei that are critical for further self-assembly [20]. This critical nucleus is defined as the smallest assembly size that grows faster by the addition of monomers, than dissociates back to smaller assemblies including monomers[21]. Once the critical nucleus has been formed, there is a rapid formation of fibrils by the addition of monomers. This is the elongation phase and fibril formation in this way is known as primary nucleation. The formation of these nuclei is therefore crucial in the generation of amyloid and the identification of these structures will ultimately aid our understanding of amyloid assembly and pathology.

One likely nucleus of Aβ42 assembly has been suggested to be a hexameric assembly[20,22-26]. It has been shown that the formation of hexameric Aβ42 is an early event in the self-assembly pathway[23,24,27,28] and the identification of several multimers of hexamers e.g. Aβ derived diffusible ligands (ADDLs), Aβ*56 and globulomers provide a compelling argument that the hexamer is the basic building block for the formation of toxic oligomers[15,29]. The majority of these studies investigating the role of hexameric Aβ42 as a nucleus for self-assembly make use of synthetic peptides which are greatly advantageous due to being readily available at high concentrations necessary for biophysical and structural characterisation. However, the true cellular environment and processing of C99 to release Aβ cannot be mimicked using these synthetic peptides.

Furthermore, familial AD (FAD) causing mutations within the Aβ sequence, which are also in the extracellular domain of the C99 sequence, have been shown to have a higher aggregation propensity in previous studies using synthetic and recombinant proteins[30-38]. It is not yet understood whether these FAD mutations which promote self-assembly, increase overall Aβ production, and/or change biochemical properties, also promote/enhance the formation of hexameric Aβ assemblies.

Here, by transfection of Chinese Hamster Ovarian (CHO) cells, we identify the formation of hexamers in Aβ enriched conditions. We also identify for the first time, the formation of hexameric Aβ in CHO transfected with the Flemish (A21G), Dutch (E22Q), Italian (E22K), Arctic (E22G) and Iowa (D23N) FAD causing mutations[39-42]. We have isolated cell-derived hexameric Aβ assemblies and assessed their assembly and nucleating properties using complementary techniques including Mass Spectroscopy, Thioflavin T (ThT) fluorescence and immunoblotting. We identify these cell-derived Aβ hexamers as Aβ42 assemblies which are major contributing nuclei for the self-assembly of Aβ monomers. This nucleating propensity is much more pronounced on monomeric Aβ42 than Aβ40 and is highly dependent on the concentration of available monomers. Furthermore, we show for the first time in a cellular context that the formation of this hexamer is an inherent property of the Aβ42 peptide and its self-assembly propensity, as a self-assembly impaired primary sequence variant of Aβ42 does not form hexamers. The identification of assemblies nucleating Aβ self-assembly in this way provides potential therapeutic targets to prevent oligomer-induced neurotoxicity.

## Results

### Aβ assembly profile in CHO cells: Identification of hexameric Aβ

Although Aβ self-assembly has been extensively studied with synthetic peptides, less is known about the assembly of Aβ peptides produced in a cellular context. As C99 processing precedes Aβ release in physiology, we first assessed the Aβ assembly profile in CHO cells transfected with the human C99 sequence affixed with the signal peptide of the full-length APP in a pSVK3 plasmid backbone. Cell lysates and media of CHO cells were harvested 48 hours after transfection and the Aβ profile was assessed by western blotting and detected using the monoclonal human specific anti-Aβ W0-2 antibody (Figure 1A). In line with what has previously been shown by our group, we confirm that cells transfected with C99 produce a detectable band corresponding to an Aβ assembly size of ∼28kDa; the theoretical size of an Aβ42 hexamer [43]. Detection with the anti-Cter antibody against the C-terminal of APP did not identify these bands, consolidating these assemblies are likely to be Aβ and do not contain the CTF of C99 (Supplemental Figure S1). Furthermore, as the band was not detected in the Empty Plasmid (EP) condition, we are confident that the hexamer is not a product of the transfection protocol. Whilst only Aβ hexamers were detected in the cell lysates of these cells, monomers, dimers, trimers and hexamers were detected in the media confirming that in our cellular model, C99 is cleaved to release Aβ detectable both intra- and extracellularly (Figure 1A). Additionally, we also assessed the Aβ profile of cells transfected with the Aβ40 and Aβ42 sequences affixed with the APP signal peptide (referred to as C40 and C42 respectively) to understand whether the formation of this hexamer in a cellular context was related to one or both of these Aβ isoforms (Figure 1B). We do not identify a hexameric band in the cell lysates or culture media of C40 transfected CHO cells, which would be ∼26kDa in size, in line with the preferential formation of hexamers by Aβ42 previously shown with synthetic peptides[23]. The identification of a hexameric band both intra- and extracellularly in C42 transfected CHO cells suggests that 1) formation of hexameric Aβ is irrespective of whether there is first the processing of the C99 fragment and 2) cell-derived hexamer formation is likely to be an intrinsic property of the Aβ42 sequence itself, as has been demonstrated previously with synthetic peptides[23,26,27].

**Figure 1.**
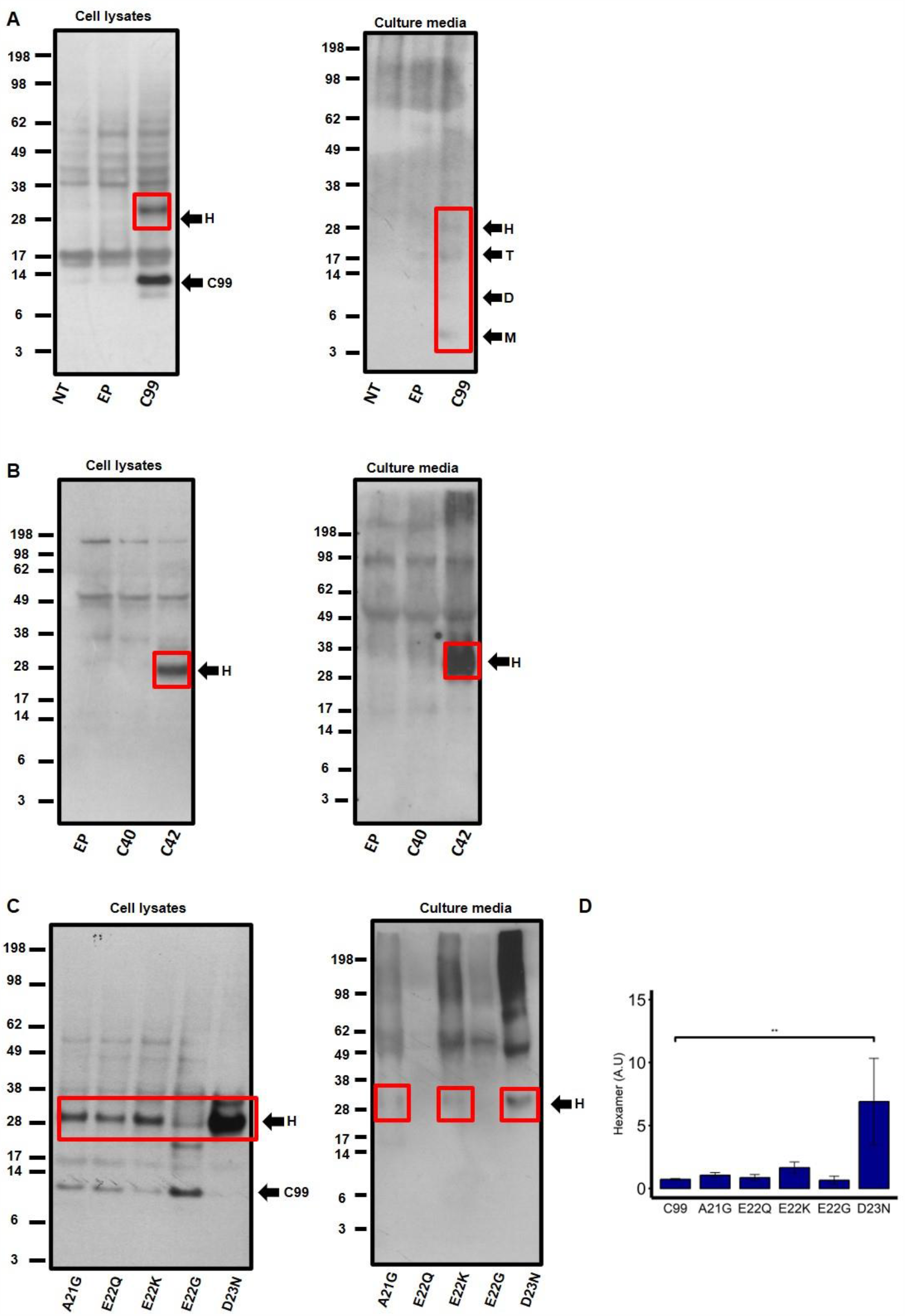
Aβ profile in transfected CHO cells detected with the anti-Aβ W0-2 antibody. (A) Hexamer formation (red box) is seen in cell lysates of CHO cells transfected with C99 (right panel). Monomers (M), Dimers (D), Trimers (T) and Hexamers (H), were detected in the culture media of C99 transfected CHO cells (left panel). (B, right panel) No assemblies were detected in the cell lysates of C40 transfected CHO cells, however, hexameric Aβ was detected in cell lysates of C42 transfected CHO cells. (B, left panel) Hexameric Aβ is also detected in culture media of CHO cells transfected with C42 only. (C) Hexameric Aβ is detected in cell lysates and media of CHO cells transfected with FAD mutations in the C99 sequence. (D) Hexamer formation was quantified on the C99 signal (black arrow) for each of the mutations. The D23N (Iowa) mutation showed a significant increase in hexamer formation (**) compared to C99. One-way ANOVA with Tukey’s post hoc comparison where p= < 0.01 (*), < 0.001 (**), < 0 (***). Error bars are expressed as ±SEM (N=3).

To link the formation of hexameric Aβ assembly to disease related conditions, we investigated the Aβ profile in CHO cells transfected with the C99 sequence containing the A21G, E22Q, E22K, E22G and D23N FAD mutations. In Figure 1C (left panel), we show that in CHO cells transfected with the C99 sequence containing these mutations, there is the detection of hexameric Aβ in cell lysates. The hexamer is also detected in the media of A21G, E22K and D23N expressing cells (Figure 1C, right panel). In particular, the D23N mutation shows a much more pronounced hexamer formation compared to any other mutant. This was confirmed by quantification of hexamer production in cell lysates normalised to the C99 signal (Figure 1D) which shows that the D23N mutant generates significantly more hexameric Aβ than the wild-type C99 (p=<0.001), while all other mutants displayed similar levels of hexamer formation. Together, this demonstrates that formation of hexameric Aβ is occurring in several AD-related conditions.

From these results, we can conclude that the formation of a hexameric assembly is a common feature in Aβ enriched conditions as well as in AD related Aβ mutations where amyloid formation is accelerated.

### Isolation and composition of cell-derived Aβ

As the aim of this study was to characterise the formation, self-assembly and nucleating properties of the hexamer, we next optimised the isolation of media derived hexameric Aβ. We have focussed here on Aβ material in the media as we hypothesise that the self-assembly leading to the deposition of extracellular amyloid plaques is likely dependent on the presence of this hexamer in the extracellular space. To isolate the Aβ hexamer, CHO cells were transfected with C42 or C99 for 48 hours and media was immunoprecipitated using the W0-2 antibody. This was then separated using the Gel Eluted Liquid Fraction Entrapment Electrophoresis (GELFrEE^®^) 8100 system. Briefly, as with SDS-PAGE, this system separates the peptide by size with the added advantage of collecting the assembly size of interest as a liquid fraction. Shown in Figure 2A, we confirm by western blotting the isolation of hexameric Aβ from the media of CHO cells transfected with C42 and C99 in fraction 5 only (C42 fraction 1-4 shown in Supplemental Figure S2). By isolating and characterising hexamers from both conditions, we are able to assess whether processing affects the self-assembly and nucleating properties of Aβ hexamers. We are also able to isolate C99-derived Aβ monomers in fraction 1, however, as lower molecular weight assemblies were not detectable in CHO cells transfected with C42, even when samples were harvested at earlier time points after transfection (Supplemental Figure S3), this was not possible for C42 transfected CHO cells. Considering the highly hydrophobic and aggregation prone nature of the Aβ42 sequence, it is unsurprising that we cannot detect lower molecular weight assemblies which, if present, are likely too low in concentration to detect by western blotting.

**Figure 2.**
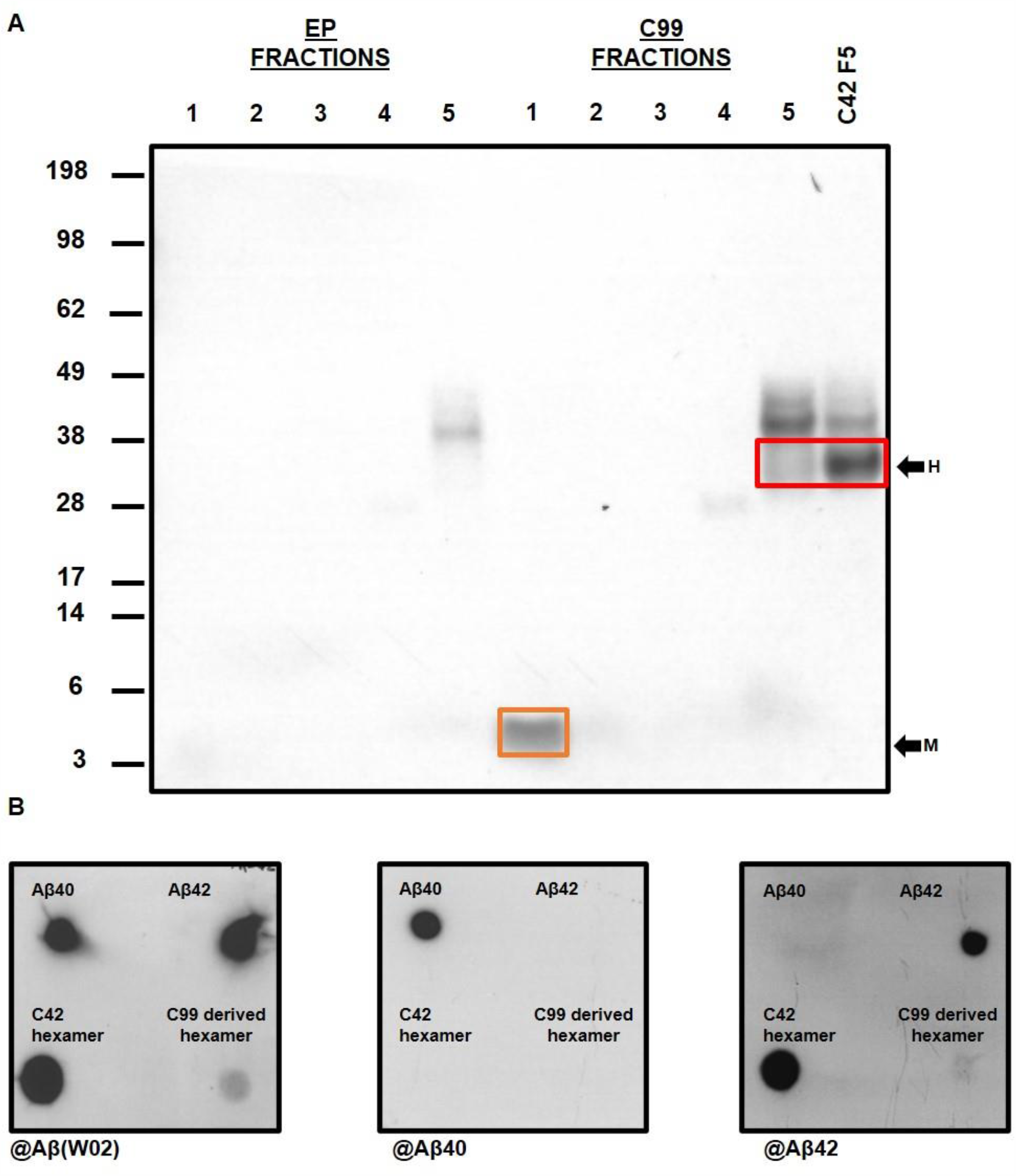
Isolation of Aβ assemblies and identification of hexameric assembly Aβ isoform. (A) Aβ was immunoprecipitated using the W0-2 antibody and separated by size using the GELFrEE^®^ 8100 system. CHO cells transfected with the empty plasmid (EP) and immunoprecipitated for Aβ did not show isolated assemblies in any fraction. From C99 transfected CHO cells, we are able to isolate monomeric Aβ (orange box) in Fraction 1 and hexameric Aβ (red box) in Fraction 5. Only hexameric Aβ was isolated (Fraction 5) from C42 transfected CHO cells. (B) Dot blotting was carried out to identify the hexameric Aβ isoform using the W0-2, anti-Aβ40 and anti-Aβ42 antibodies. Synthetic Aβ40 and Aβ42 were used as positive controls. Both C42 and C99 derived Aβ hexamers are detected by the anti-Aβ W0-2 antibody (left panel) and anti-Aβ42 antibody (right panel), however, not by the anti-Aβ40 antibody (middle panel) confirming the hexamers are composed of Aβ42.

To identify the isoform of Aβ in the C42 and C99-derived hexameric assemblies, we carried out dot blotting using anti-Aβ42 and anti-Aβ40 specific antibodies (Figure 2B) with synthetic preparations of Aβ40 and Aβ42 as positive controls. Dot blotting with W0-2 (Figure 2B, left panel) antibody was used to confirm the presence of the proteins. Our results clearly show the Aβ42 specific antibody binds to hexameric assemblies from both conditions (Figure 2B, right panel) whereas no signal is seen for either hexamer with the Aβ40 specific antibody (Figure 2B, middle panel), thus confirming the Aβ hexamers are Aβ42 assemblies.

### The formation of a hexameric assembly is an inherent property of the Aβ42 peptide due to self-assembly propensity

The preferential formation of hexameric Aβ has been suggested to be linked to the C-terminus of Aβ42 and its self-assembly propensity[23]. To assess this in our cellular model, CHO cells were transfected with an assembly-impaired variant of the C42 sequence, vC42, and the Aβ profile was assessed by western blotting and detection with the W0-2 antibody (full primary sequence can be found in Supplemental Table S1). This is the same sequence that has been previously reported and thoroughly characterised as being self-assembly impaired despite only a two amino acid difference (F19S and G37D) compared to the wild-type Aβ42 sequence[44]. Figure 3 demonstrates that CHO cells transfected with this variant produced only dimers in the cell lysates and no detectable assemblies in the media. The lack of a hexameric assembly confirms the importance of self-assembly in the formation of this structure and its direct link to Aβ42 aggregation propensity.

**Figure 3.**
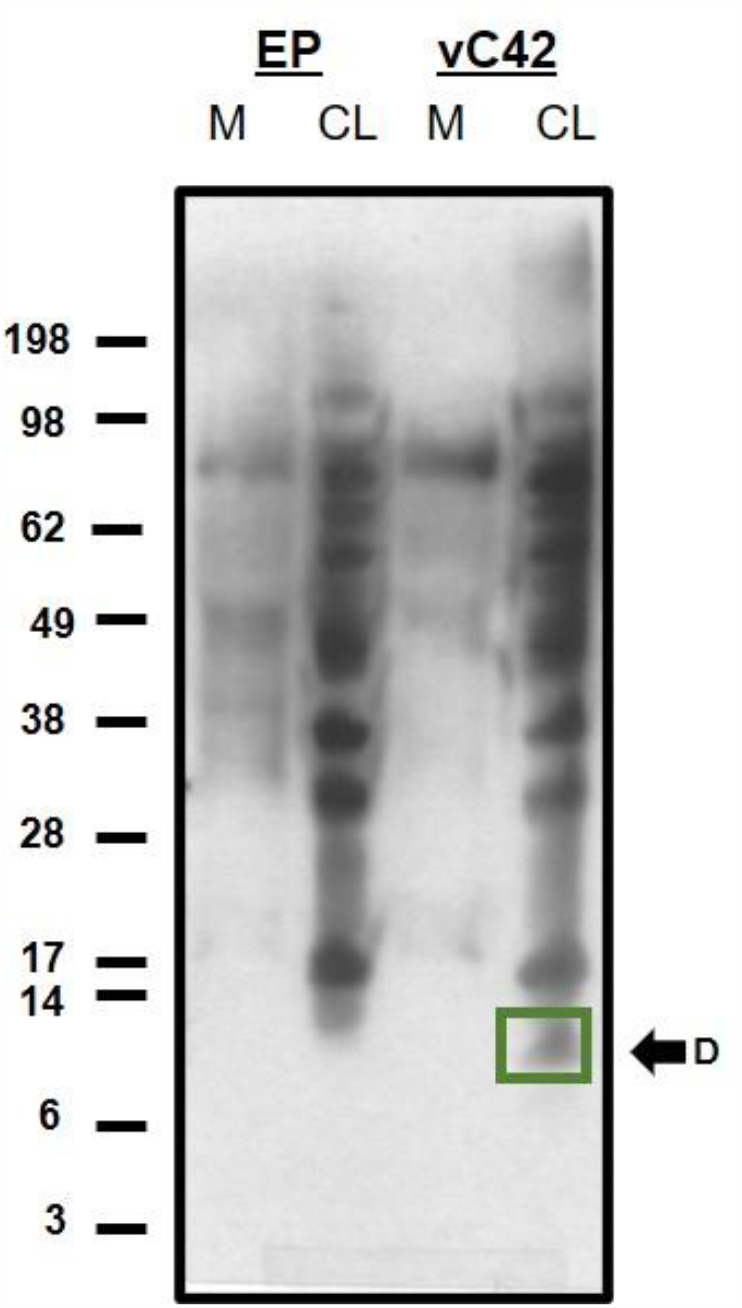
Hexamer formation is directly related to Aβ42 self-assembly propensity. CHO cells transfected with vC42 (F19S, G37D), did not form hexameric Aβ in either the cell lysates or media detected by the W0-2 antibody. However, dimers (green box) were detected in the cell lysates of vC42 transfected CHO cells.

### Cell-derived Aβ42 hexamer formation is a direct consequence of Aβ42 primary sequence

As both monomeric and hexameric Aβ were detected and isolated from C99 transfected cells, we next questioned whether the C99-derived Aβ monomers isolated in Fraction 1 (Figure 2A) were able to assemble into hexamers. No assembly of the Aβ monomer isolated from the media of the C99 transfected cells was seen by western blotting over 48hours, the time in which we see hexamer presence in the cell lysates and media of C42 and C99 transfected CHO cells (Figure 4A). Electro-chemiluminescence immunoassay (ECLIA) measurements, which provide quantitative analysis of monomeric Aβ, were performed on the media of C99 transfected CHO cells (Figure 4B) before immunoprecipitation for Aβ and confirmed that Aβ40 was in high abundance (80.9pg/mL) whereas very low concentrations of Aβ42 (1.3pg/mL) were detected. This provided early indications that the monomer detected by western blot (Figure 1A) is likely to be Aβ40. Analysis of the isolated monomer fraction by mass spectrometry (Figure 4C) revealed that, in line with the reduced self-assembly properties reported in the literature[45], only the Aβ40 sequence was identified in this sample. Finally, we have confirmed by western blotting that Aβ40 is unable to form hexamers even in a cellular context, as CHO cells transfected with C40 do not produce a detectable band for this assembly in either cell lysates or media (Figure 1B). qPCR data consolidated that this was not due to low transfection efficiency (Supplemental S4).

**Figure 4.**
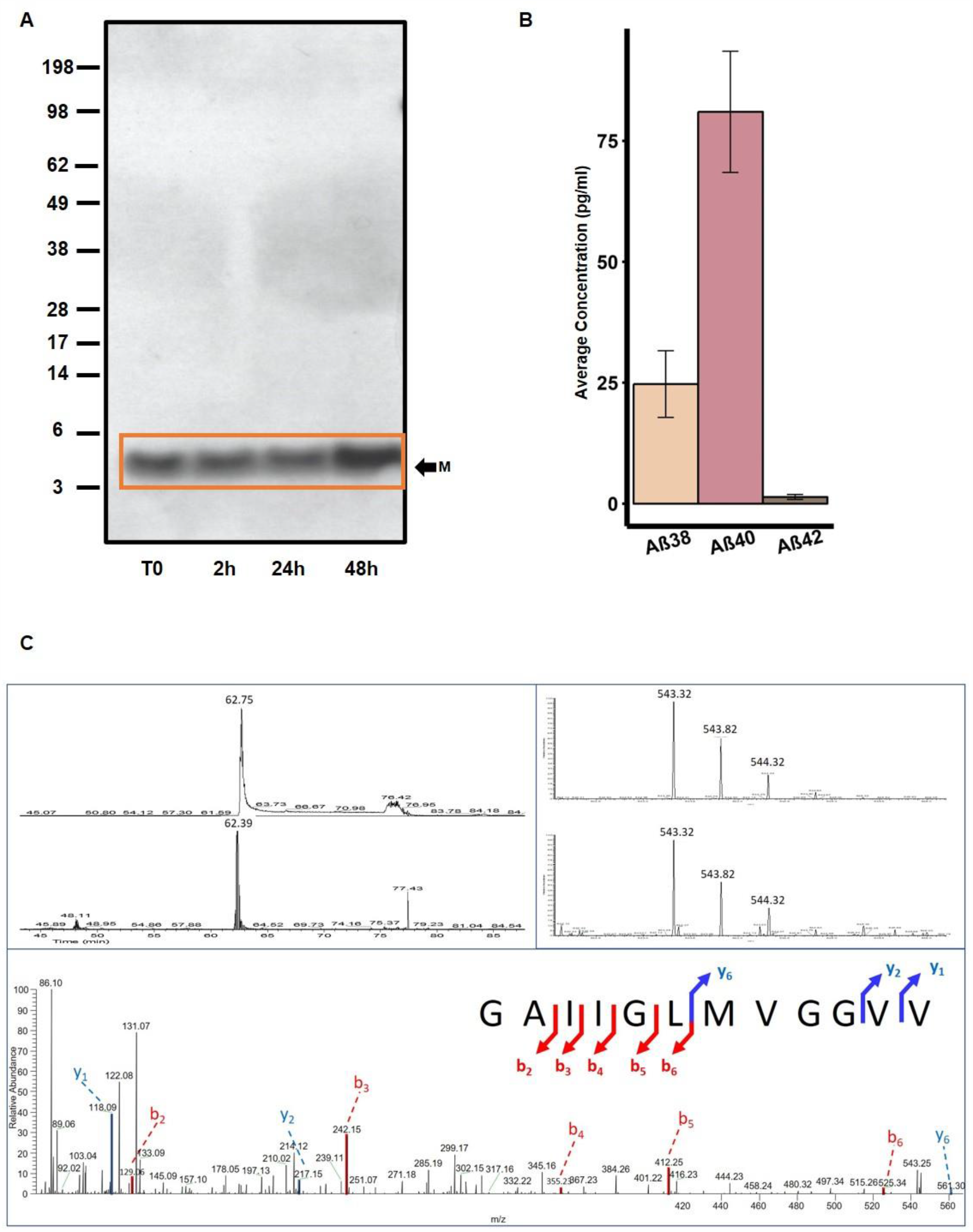
Hexamer formation is directly related to Aβ42 primary sequence. (A) Isolated monomeric Aβ incubated at room temperature over 48hrs does not assemble into hexameric or other higher molecular weight assemblies as assessed by western blotting detected with the W0-2 antibody. (B) ECLIA measurements (N=5) on the media of C99 transfected CHO cells identified Aβ40 as the most abundant monomeric Aβ isoform (80.9pg/ml). Low concentrations of Aβ38 (24.7pg/ml) and even lower concentrations of Aβ42 (1.3pg/ml) were detected in this culture media (C) CLC-MS/MS identification of Aβ40 in the C99 Sample. Top left panel: extracted ion chromatogram of the [M+2H]2+ detected at 62 minutes, for the standard (top) and the C99 sample (bottom). Top right Panel: mass spectra of the Aβ40 fragment ion showing the isotopic pattern for the standard (top) and the C99 sample (bottom). Bottom panel: MS/MS annotated spectrum characterising an Aβ40 fragment from the C99 sample.

Together, these data show for the first time from cell-derived material that Aβ40 does not readily form hexameric assemblies, which in turn reinforces that the primary sequence of Aβ42 and its resultant self-assembly properties are determining factors in the formation of a hexameric assembly.

### Isolated hexameric Aβ42 does not self-assemble into higher molecular weight assemblies

In order to establish the self-assembly properties of the isolated hexameric Aβ42 derived from media of C42 and C99 transfected CHO cells (Fraction 5 shown in Figure 2A), we first carried out a ThT fluorescence assay as a measure of fibril formation. 150µM of hexameric Aβ was incubated with 20µM ThT and fluorescence was monitored over 48 hours. Figure 5A shows that over this time course, there is no increase in fluorescence seen which suggests that the hexameric assembly does not form fibrils in the timeframe of our experiment. To further consolidate this and detect any pre-fibrillar assemblies that may not bind to the ThT dye, western blotting was carried out on the same samples at several time points (Figure 5B and C). Detected by the W0-2 antibody, we see there are no higher molecular weight assemblies at longer time points and only the hexameric assembly is detected for C42-derived hexamers. This also confirms that there is no degradation or disassembly of the peptide. However, although C99-derived Aβ42 hexamers do not assemble into higher molecular weight assemblies, monomers are detected with increasing intensities over time. This suggests that in the C99 transfected conditions where processing is taking place, the hexamer is less stable and does disassemble into monomers. This is reflective of the dynamic nature of self-assembly, particularly in the formation of a critical nucleus. As we have identified the hexamers to be only composed of the Aβ42 isoform, these data also allow us to conclude that the monomers detected from the disassembled C99-derived hexamers are likely to be Aβ42 monomers. Combined, these data show no further self-assembly of hexameric Aβ assemblies derived from both C42 and C99 in our experimental conditions. We therefore conclude that these cell-derived hexameric assemblies have similar self-assembly properties in our experimental conditions; the hexamer on its own does not self-assemble into higher molecular weight assemblies, however, the stability of C99-derived Aβ42 hexamers is weaker than that of C42-derived Aβ42 hexamers.

**Figure 5.**
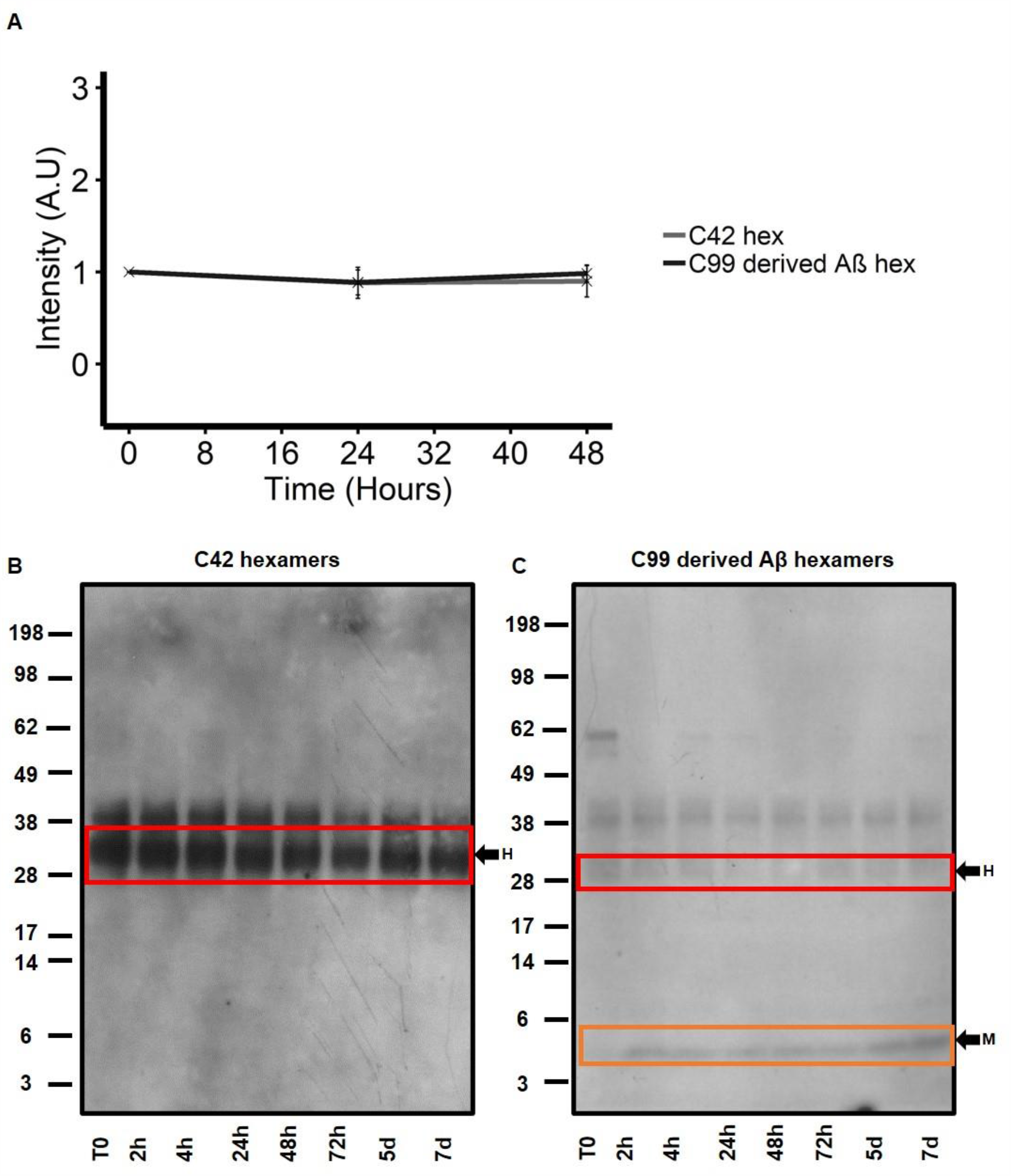
Isolated hexameric Aβ does not self-assemble into higher molecular weight assemblies. (A) 150µM isolated hexameric Aβ from C42 and C99 transfected CHO cells was incubated with 20µM ThT and fluorescence was monitored over 48hrs as a measure of fibrillogenesis. No increase in fluorescence was seen for either of the isolated hexamers. (B and C) Western blotting and detection with the W0-2-antibody revealed both hexamers (red box) do not assemble into higher molecular weight assemblies over 7days. Hexamers from C99 transfected CHO cells, do, however disassemble into monomers (orange box) over time as was detected from 2hrs onwards.

### Hexameric Aβ42 preferentially nucleates self-assembly of Aβ42 monomers

Finally, we assessed the nucleating properties of isolated Aβ42 hexamers by seeding monomeric synthetic Aβ42 (mAβ42) with increasing amounts of isolated C42 and C99-derived Aβ42 hexamers (Figure 6A). mAβ42 was prepared as previously described [44] and diluted to a working stock concentration of 50µM. ThT fluorescence of mAβ42 without any seeding (Supplemental Figure S5) shows a lag phase (0-4hours) and an elongation phase (4-24hours), as has been previously shown[44,46,47]. The concentration of hexamer used for seeding was a percentage of the mAβ42 concentration and the final solution was incubated with 20µM ThT dye. As the nucleating effects of the hexamer are expected to be in the early stages of assembly, ThT fluorescence was monitored for 4 hours and normalised for each condition to itself at T0 as a representation of increased fluorescence at each time point (Figure 6A).

**Figure 6.**
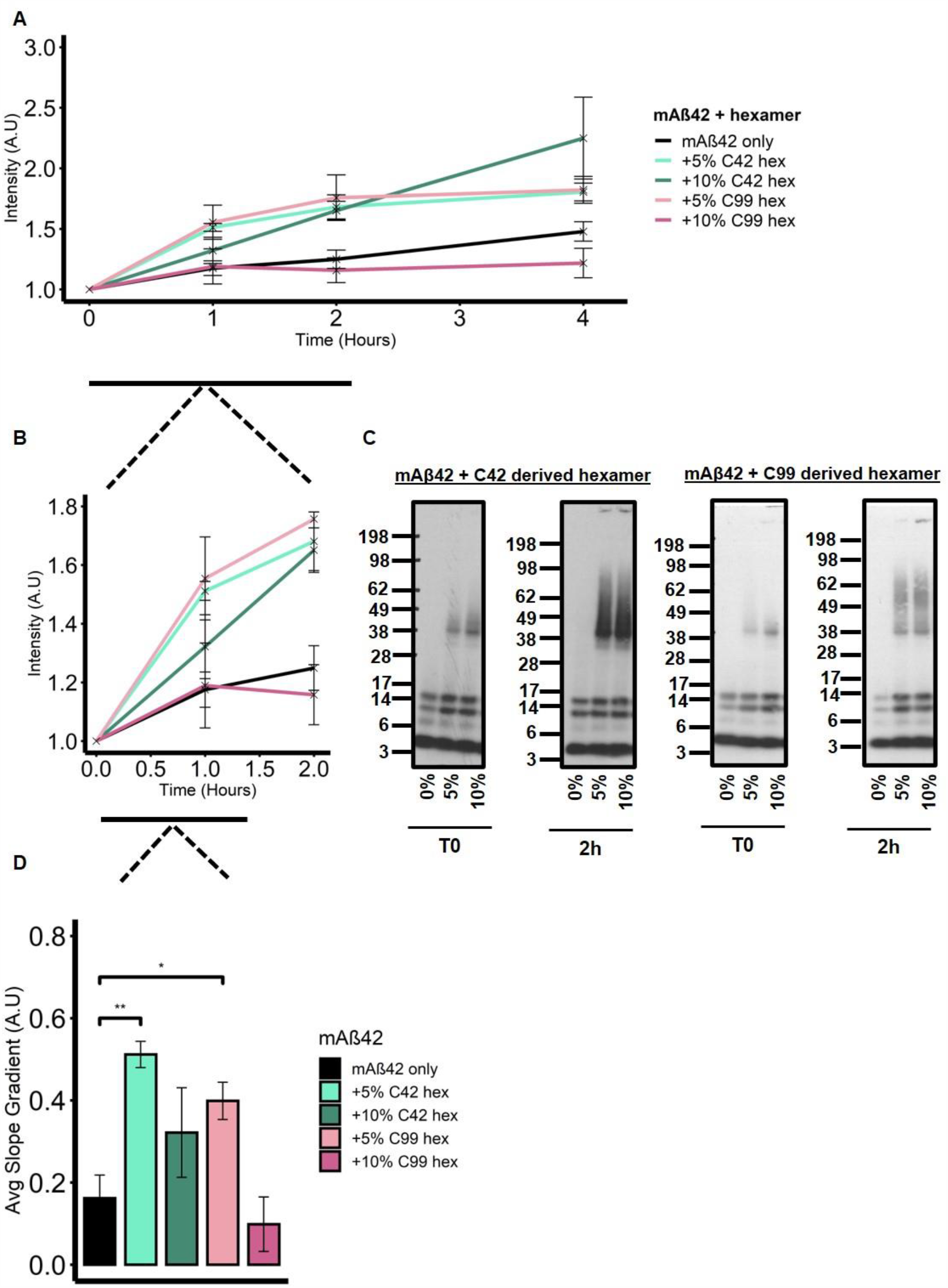
Isolated C42 and C99 derived hexameric Aβ nucleate monomeric Aβ42 (mAβ42). (A) 50µM mAβ42 was seeded with 5- or 10% hexamer and the solution was the incubated with 20µM ThT. Fluorescence was monitored over 24 hours. (B) Addition of both 5- and 10% C42 hexamer results in an immediate increase in ThT fluorescence (0-2 hours). (C) Western blotting with the W0-2 antibody revealed all seeding conditions form higher molecular weight assemblies at T0 and by 2hrs fibrils are seen ‘stuck’ in the wells of the gel. (D) The gradient of the ThT fluorescence slope was calculated at the early time points of aggregation (0-1 hour) and One-way ANOVA with Tukey’s post hoc comparison where p= < 0.01 (*), < 0.001 (**), < 0 (***), revealed addition of 5% C42 hexamer significantly (**) increased the kinetics of aggregation compared to mAβ42 only as did the addition of 5% C99 derived hexamer (*). Error bars are expressed as ±SEM.

The addition of 5% (light pink line) and 10% (dark pink line) C42-derived hexamer immediately results in an increase in fluorescence intensity, as does the addition of 5% (light green line) C99-derived Aβ42 hexamer. This suggests increased self-assembly kinetics in the early stages of aggregation for these conditions, however, 10% (dark green line) addition of C99-derived Aβ42 hexamer results in a similar ThT fluorescence as mAβ42 alone (black line). This was further consolidated by assessing the range of assembly sizes present at T0 and 2 hours after hexamer addition by western blotting using the W0-2 antibody (Figure 6C). We see that even at T0, mAβ42 seeded with 5 and 10% C42-derived hexamers show the formation of higher molecular weight assemblies which are not present in the mAβ42 only sample. By 2 hours, both seeded conditions show the formation of a larger range of higher molecular weight assemblies which migrate as a smear as well as fibrils ‘stuck’ in the well of the gel, whilst mAβ42 without seeding displays bands corresponding to monomers, dimers and trimers only. Western blotting for mAβ42 seeded with C99-derived Aβ42 hexamers (Figure 6C) also revealed similar trends to that of C42-derived hexamer seeding where higher molecular weight assemblies were detected at T0 in the seeded conditions, and by 2hours fibrils were ‘stuck’ in the wells of the gel. As we have identified both hexamers to be Aβ42, this is unsurprising. Interestingly, in contrast to what was seen with the ThT fluorescence, 10% addition of C99-derived Aβ42 hexamers does result in the formation of higher molecular weight assemblies and fibrils being stuck in the well by 2 hours, perhaps suggesting the formation of ThT negative aggregates. Combined, this supports the hypothesis of the hexamer as a nucleus for self-assembly.

For a more robust analysis of the nucleating effects in the early stages of self-assembly, we have calculated the gradient of the graph for each condition from 0-1 hour as an indication of assembly kinetics (Figure 6D). The addition of 5% C42 and C99-derived hexamer significantly increases the gradient of the graph (0.51 AU ± 0.03 SEM, 0.4AU ± 0.05 SEM respectively, p= <0.01) compared to mAβ42 alone (0.16 AU ±0.05 SEM). Although an increase was also seen with 10% C42 hexamer (0.32 AU ± 0.1 SEM), this was not significant. This is likely due to the fact that the nucleating potential of the cell-derived C42 hexamer is dependent on the available monomers in solution. Furthermore, the same increase in early assembly kinetics was not seen with 10% C99-derived Aβ42 seeding (0.1 AU ±0.06 SEM). This might be due to the difference in stability of the C99-derived Aβ hexamer; as it disassembles into monomers with time (Figure 4C), the concentration of monomers continues to dominate the solution population and there is perhaps not enough hexamer in solution to nucleate self-assembly which may indicate a threshold concentration is required before nucleating effects can be seen.

Finally, we also assessed the nucleating properties of both hexamers on monomeric Aβ40 (mAβ40) prepared using the same protocol as for mAβ42 (Figure 7). No increase in slope gradient at early time points (0-1 hour) was seen for mAβ40 seeded with 5% and 10% C42 or C99-derived Aβ42 hexamers respectively. The lack of nucleating effects at early stages of self-assembly was further consolidated by western blotting (Figure 7C) which revealed assembly sizes ranging from monomers to tetramers only for all conditions at T0 and 2 hours and no increase in higher molecular weight species. The ThT fluorescence for both 5 and 10% C42-derived hexamer seeding does begin to slightly increase after 4 hours which could be indicative of a reduced ability of these hexamers to nucleate Aβ40 compared to Aβ42. A similar and more pronounced trend of increased ThT fluorescence is seen with 5 and 10% C99-derived Aβ hexamers from 2 hours onwards. Interestingly, as the increase in fluorescence was not seen in mAβ42 seeded with 10% C42-derived hexamers (Figure 6) and as we have shown the C99-derived hexamers to disassemble into monomers, this data suggests some effect of two monomeric Aβ isoforms interacting.

**Figure 7.**
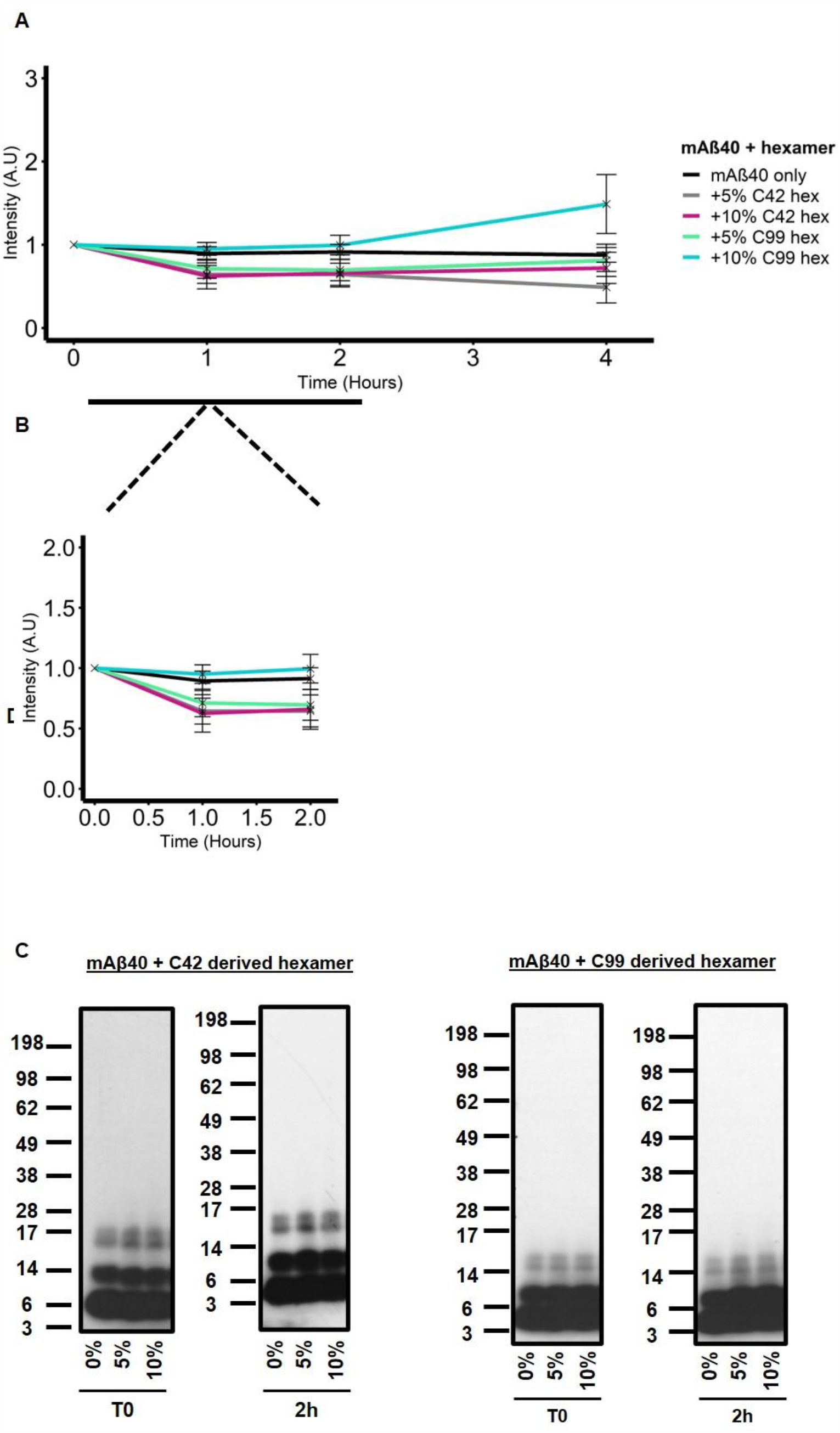
Isolated C42 and C99 derived hexameric Aβ do not readily nucleate monomeric Aβ40 (mAβ40). (A) 50µM mAβ40 was seeded with 5- or 10% hexamer and the solution was then incubated with 20µM ThT. Fluorescence was monitored over 4 hours. (B) Addition of both 5- and 10% did not result in an increased fluorescence at early time points (0-2 hourss) (C) Western blotting detected with the W0-2 antibody revealed seeding conditions with both hexamers does not lead to the formation of higher molecular weight assemblies compared to mAβ40 only.

Together, we conclude that the cell-derived Aβ42 hexamers have a reduced nucleating propensity on Aβ40 which further reiterates the direct link of hexamers as critical nuclei for Aβ42 self-assembly. To be sure of this reduced capacity as opposed to inability, we seeded mAβ40 with 30% C42 and C99-derived hexamers (Supplemental Figure S6), which confirms that with enough hexamer, seeding can occur.

Our data demonstrate for the first time, the ability of cell-derived Aβ42 hexamers to nucleate self-assembly with preferential nucleation of monomeric Aβ42 compared to Aβ40.

## Discussion

The process of self-assembly and its importance in Aβ toxicity has been the focus of several research studies. The formation of intermediary oligomeric species during this process has been identified as being a determinant of cytotoxicity and we therefore investigated an Aβ assembly that is responsible for facilitating nucleation dependent amyloid formation. To the best of our knowledge, we are the first group to present a thorough characterisation of cell-derived hexameric Aβ and provide more physiologically relevant evidence than *in vitro* studies using synthetic or recombinant peptides to further support this assembly to be a nucleation enhancing entity.

The identification of specific Aβ intermediate assemblies that serve as nuclei for fibril formation has remained elusive due to their transient and heterogenous nature. Despite this, several studies have optimised the use of highly sensitive techniques such as small angle neutron scattering (SANS), small angle x-ray scattering (SAX) and sedimentation velocity (SV) analysis complementary to SDS-PAGE of photo-induced cross linking of unmodified proteins (PICUP) solutions of Aβ to detect hexameric assemblies involved in the early stages of self-assembly[26-28]. Furthermore, a recent native ion mobility-mass spectrometry study has also identified the formation of hexameric Aβ and suggests a β-barrel structure in membrane mimicking environments[48]. In line with our data, these studies have all consistently observed the formation of hexameric Aβ to be highly prone to the Aβ42 sequence. However, these studies have relied heavily on synthetic peptides which cannot mimic a cellular environment. Here, in our experimental conditions, we have identified a non-self-assembling Aβ42 specific hexamer that is present in both the cell lysates and media of transfected CHO cells.

CHO cells transfected with either C99 or C42 sequences showed the ability to form an Aβ assembly that was ∼28kDa in size by western blotting, which is the theoretical size of an Aβ42 hexamer. FAD Aβ mutations in the C99 sequence also showed the formation of hexameric Aβ in the cell lysates and media of the CHO cells suggesting that the formation of this assembly is common in Aβ enriched and FAD related conditions. The commonality of hexameric Aβ across these conditions highlights for the first time in a more physiological context, the importance of this assembly in conditions where Aβ self-assembly is accelerated.

Dot blotting following the isolation of these hexamers from the media of C42 and C99 transfected cells, confirmed them to be Aβ42 assemblies (Figure 2B). On the contrary, the monomeric Aβ identified and isolated from the media of C99 transfected CHO cells was confirmed to be composed of the Aβ40 sequence only. Furthermore, this monomer did not assemble into hexamers or any other higher molecular weight assemblies in the parameters of our experiments. This information is important as it confirms that the ability to readily form a hexameric assembly in a cellular context is an inherent property of the Aβ42 primary sequence. This was further supported by the lack of a hexameric assembly seen in the cell lysates and media of CHO cells transfected with the vC42 sequence which has both F19S and G37D substitutions. These substitutions have been shown to negatively affect self-assembly propensity[35,49-55]. In this way, vC42 also begins to provide some evidence to suggest that both the F19 and G37 amino acids are important residues in the formation of a hexameric assembly. Interestingly, whilst this peptide was shown to remain largely monomeric for 7 days *in vitro*[44] we show here in cellular context, the formation of dimers within 48 hours. This further highlights the physiological relevance of our study in which we show the importance of translation of the protein, the cellular environment and the subsequent Aβ42 assemblies formed.

Our data strongly supports the conclusion that the Aβ hexamers we have identified are Aβ42 specific assemblies (Figure 2B). This, combined with the assumption that there are no other isoforms present in the C42 condition, as well as C99-derived hexameric Aβ disassembling back into monomers over time (Figure 5C), suggests the initial nascent Aβ42 monomer may be responsible for the formation of the Aβ42 hexameric structures identified here. A previous study exploring the decapeptide Aβ (21-30) showed its protease resistance was identical to full length of Aβ42 and likely to organise intramolecular monomer folding and therefore an initial folding nucleus[56]. This has been attributed to a β-turn formed and stabilised by both hydrophobic and electrostatic interactions between V24-K28 and K28-E22/D23. FAD related mutations at positions G22 or D23 therefore disrupt this turn stability and have been shown to enhance subsequent assembly and oligomerisation[54,57,58]. The level of turn disruption correlates directly with enhanced oligomerisation for each mutation; from our results, the D23N mutation significantly disrupts the turn stability and enhances the formation of hexameric Aβ42. The importance of this monomer folding nucleus in the formation of higher molecular weight assemblies, such as the hexamer, has been explained by the formation of the stabilised turn being a kinetically favoured folding event capable of facilitating the interaction between the central hydrophobic cluster (L17-A21) and the C-terminus, which is far more pronounced in Aβ42 than in Aβ40[58]. Together, this provides a plausible explanation as to 1) how the hexameric assembly is linked to folding events in the monomeric Aβ42 peptide 2) why FAD mutations explored in this study do not negatively affect hexamer formation[59]. The disruption of the stabilising interactions in the decapeptide region of monomeric Aβ may occur in a physiological environment such an acidic pH (e.g. endosomes) where Aβ assembly is known to be enhanced.

We also show that hexameric assemblies that are formed from Aβ after C99 processing are less stable than those formed from C42 where there is no upstream processing, suggesting processing may have an effect on structural properties. Despite this, both hexamers display nucleating properties which points to size being an important contributor for nucleating potential. Interestingly, hexamers have also been identified as being important intermediates in the self-assembly of β2-microglobulin which suggests this assembly size may play an important role in the aggregation of several amyloid forming proteins[60].

The lack of self-assembly into higher molecular weight assemblies (Figure 5) over 7 days suggests that the hexamer we have identified may be an ‘off-pathway’ oligomer. However, several oligomeric species that are multimers of a hexameric unit e.g. ADDLs and Aβ*56 which would likely require hexameric self-association have been identified. It is beyond the scope of this study to firmly identify the Aβ hexamer presented here as an off-pathway oligomer, however, in the parameters of our experiments this may be the case. Hexamers with the ability to self-associate may be in a different conformation and/or require suitable conditions for self-association such as membrane interactions or an acidic environment. Furthermore, the lack of association of two hexamers to form a dodecamer, which is thought to occur due to stacking of two hexamers with their hydrophobic C-terminal ends at the centre of the structure[22] was also observed by Osterlund and colleagues. They concluded the reduced entropic drive towards hexamer dimerisation is likely due to the C-termini being stabilised in their experimental conditions which are likely mimicking the effects that would be seen in a lipid bilayer [48]. Therefore, perhaps the hexamers we isolate here are or have been associated to these lipid bilayers which affects their self-association properties.

Importantly, we show for the first time the nucleating potential of both C42 and C99 cell-derived hexamers and show this to be preferential to mAβ42 over mAβ40. This nucleation propensity is heavily reliant on the available monomers in solution, in line with the definition of a nucleus for self-assembly during the lag phase of amyloid formation[21]. We believe that as the hexamer is not a fibrillar species, it is likely directly involved in primary nucleation especially due to kinetics of assembly increasing at the very early time points of assembly when secondary nucleation, where newly formed fibrils provide the surface to catalyse new aggregates from the available monomers, is likely not yet occurring. The switch to secondary nucleation could occur as early as 2 hours as this is when we see the formation of fibrils in the well of the gels by western blotting. Secondary nucleation is dependent on both the concentration of monomers and existing fibrils, and as the gradient of the slope suggests slower aggregation of mAβ42 seeded with 10% C42 hexamer compared to 5% between 0-1hr, perhaps there are more available monomers for secondary nucleation to occur in the later stages of self-assembly leading to more fibril formation and therefore higher ThT fluorescence[34]. However, to make solid conclusions regarding the microscopic steps involved in the self-assembly process, further in-depth work is required.

Overall, we demonstrate for the first time in a cellular context, the formation of hexameric Aβ42 as a common feature in conditions where Aβ42 self-assembly is accelerated and we have characterised these hexamers to be non-self-assembling entities that preferentially nucleate the aggregation of monomeric Aβ42. Understanding mechanisms that can enhance or facilitate self-assembly in this way will ultimately aid our understanding of amyloid pathology.

## Materials and Methods

### Chemicals and reagents

Nitrocellulose membranes were purchased from GE Healthcare (Little Chalfont, UK) and Western Lightning^®^ Plus-ECL from PerkinElmer (Waltham, MA, USA). The anti-Aβ W0-2 (MABN10) primary antibody was from Abcam (Cambridge, UK), anti-Aβ40 and anti-Aβ42 primary antibodies were purchased from Merck Millipore (Darmstadt, Germany). The anti-C-ter primary antibody and horse radish peroxidase (HRP)-conjugated secondary antibodies were purchased from Sigma-Aldrich (St Louis, MO, USA). TRIzol™ reagent and Complete™ protease inhibitor cocktail were from Roche (Basel, Switzerland). The cDNA synthesis kit and iQ SYBR Green Supermix were from Bio-Rad (Hercules, CA, USA).

### DNA constructs

The pSVK3 empty plasmid (EP) as well as the -C40, -C42 and -C99 vectors including the fused signal peptide of APP were described previously [43,61]. QuickChange^®^ site-specific mutagenesis (Stratagene, La Jolla, CA, USA) was used to produce the FAD mutants in the pSVK3-C99 template DNA, as previously described [62]. Primer sequences can be found in Supplemental Table S2.

### Cell Culture and Transfection

Chinese hamster ovary (CHO) cell lines were cultured in Ham’s F-12 medium supplemented with 10% of FBS and 1% Penicillin-Streptomycin (Life Technologies, Carlsbad, CA, USA). All cell cultures were maintained at 37°C in a humidified atmosphere (5% CO_2_). Cells were passed every 4 days at ∼80% confluency and no longer used after passage 20.

For transfections with C99 and Aβ sequences, approximately 2.2×10^6^ CHO cells were plated 24 hours in advance in 10cm petri dishes. A transfection mix of 15µg of DNA, 30µl Lipo2000^®^ (Invitrogen) in 1ml Opti-MEM® was incubated for 15mins at room temperature before being added to cells. A 0% FBS medium change was carried out 24hours after transfection and both cells and media were harvested after 48hours of initial transfection.

### Western and dot Blotting

After transfection, cell lysates were harvested and sonicated in lysis buffer (Tris 125mM pH 6.8, 4% sodium dodecyl sulfate, 20% glycerol) with Complete™ protease inhibitor cocktail. Media were centrifuged at 1200g for 5mins to pellet any debris and dead cells and the supernatants were lyophilised by SpeedVac™. For cell lysates, 40µg of protein were heated for 10mins at 70°C in loading buffer (lysis buffer supplemented with 50mM DTT and LDS sample buffer). The lyophilised media were resuspended in 400µl milli-Q (mQ) water and the maximum sample volume was loaded. Samples were loaded and separated on 4–12% NuPAGE™ bis-tris gels (Life Technologies), and then transferred for 2hours at 30V onto 0.1µm nitrocellulose membranes. After 30mins of blocking (5% non-fat milk in 0.1% PBS-Tween), membranes were incubated at 4°C overnight with primary antibodies. Membranes were then washed three times in 0.1% PBS-Tween for 10mins and incubated with horse radish peroxidase (HRP)-conjugated secondary antibodies for 1hr at room temperature. Finally, membranes were again washed three times for 10mins in PBS-Tween prior to ECL detection. Primary antibodies dilutions for western blotting are as follows: anti-Aβ W0-2 (1:1.500) and anti-C-ter (1:2.000). Secondary antibodies dilutions are as follows: anti-mouse (1:10.000) or anti-rabbit (1:10.000). Both primary and secondary antibodies were diluted in 0.1% PBS-Tween.

For dot blotting, 5µl of sample (150µM isolated Aβ hexamers, 50µM synthetic monomeric Aβ) were spotted onto 0.1µm nitrocellulose membranes. Once the samples were dry, a further 5µl of sample were spotted twice on top and dried. The membranes were boiled in PBS for 3mins twice and blocked in 5% non-fat milk in PBS-Tween for 30mins. After this the membranes were washed, incubated with antibodies and detected with ECL as described above. Primary antibodies dilutions for dot blotting are as follows: anti-Aβ W0-2 (1:1.500), anti-Aβ40 (1:1.000), anti-Aβ42 (1:1.000). Secondary antibodies dilutions are as follows: anti-mouse (1:10.000) or anti-rabbit (1:10.000). Both primary and secondary antibodies were diluted in 0.1% PBS-Tween.

### Isolation of Aβ assemblies: Gel Eluted Liquid Fraction Entrapment Electrophoresis (GELFrEE)

6.6×10^6^ CHO cells were seeded in T175 flasks 24 hours before transfection. Cells were transfected with 45µg of DNA using Lipofectamine^®^ 2000 and a 0% FBS medium change was carried out 24hours after transfection. The media were harvested and lyophilised 48hours after initial transfection and resuspended in 1ml milli-Q water. Aβ was immunoprecipitated using Sepharose A beads (Invitrogen) coated with the anti-Aβ W0-2 antibody. For immunoprecipitation, 100µl recombinant Sepharose A beads (50mg/ml) were incubated with the medium for 3hours as a pre-clearing step. This was then centrifuged at >15.000g for 5mins and the beads were discarded. The supernatant was next incubated with 5µl W0-2 antibody for 1hr at 4°C, after which 100µl fresh Sepharose beads were added and incubated at 4°C overnight. The beads were then washed 3 times with mQ water and resuspended in 104µl mQ water, 16µl of DTT 0.5M and 30µl Tris-Acetate loading buffer. The mixture was boiled at 95°C for 10mins and centrifuged at >15.000g for 5mins. The supernatant was loaded into the GELFrEE^®^ 8100 system and run using the following method; Step 1-16 mins at 50V, Step 2-40mins at 50V (Monomer fraction), Step 3-4mins at 50V, Step 4-6 mins at 70V, Step 4-13mins at 85V and Step 6-38mins at 85V (Hexamer Fraction).

Samples were collected in the system running buffer (1X buffer; 1% HEPES, 0.01% EDTA, 0.1% SDS and 0.1% Tris) and kept on ice. The monomeric fraction was then put through a buffer equilibrated 7K MWCO Zeba buffer-exchange column (Thermo Scientific) to remove the Tris-Acetate blue sample buffer. The absorbance at 280nm was read using a BioPhotometer^®^ D30 (Eppendorf) and the concentration of the collected Aβ assemblies were calculated using the molecular coefficient of 1490 M-1cm-1; (A280/1490) x1000 x 1000.

For assembly over time experiments, monomeric and hexameric samples were incubated at room temperature and 20µl aliquots were taken for western blotting at each time point.

### ECLIA

Quantification of Aβ_38_, Aβ_40_, and Aβ_42_ monomeric peptides in the serum free media of C99 transfected CHO cells was achieved using the Aβ multiplex electro-chemiluminescence immunoassay (ECLIA; Meso Scale Discovery, Gaithersburg, MD, USA) as previously described [63]. Aβ were quantified with the human Aβ specific 6E10 multiplex assay according to the manufacturer’s instructions.

### Synthetic monomeric Aβ preparation and seeding

Monomeric solutions of Aβ were prepared as previously described [44,46]. Briefly, recombinant Aβ40 and Aβ42 were purchased from rPeptide as 1,1,1,3,3,3-Hexafluoro-2-propanol (HFIP) films. 0.2mg aliquots of peptide were solubilised in 200μl HFIP (Sigma-Aldrich) to disaggregate any preformed aggregates. The solution was then vortexed for 1min and sonicated in a water bath for 5mins. The HFIP was then dried off using a steady flow of nitrogen gas. 200μl of anhydrous dimethyl sulfoxide (DMSO) (Sigma-Aldrich) was then added and the solution was vortexed for 1min. The solution was then put through a buffer equilibrated 7K MWCO Zeba buffer-exchange column (Thermo Scientific) at 4°C. The protein solution was then kept on ice whilst the absorbance at 280nm was measured using a BioPhotometer^®^ D30 (Eppendorf) spectrophotometer. The concentration was calculated using the molecular coefficient of 1490 M-1cm-1; (A280/1490) x1000 x 1000. Solutions were immediately diluted to 50μM in buffer and this was taken to be the new working stock.

### ThT Assay

To assess the self-assembly of the isolated hexameric Aβ, the fluorescence of 20μM ThT (Sigma-Aldrich) in 150μM isolated hexamer was measured over 48hours in 96 well plates using the VICTOR^®^ Multilabel Plate Reader (PerkinElmer, Waltham, MA, USA). For seeding experiments, fluorescence of 20μM ThT in 50µM monomeric Aβ40 or Aβ42 was monitored over 24hours. Fluorescence readings were obtained at room temperature with excitation and emission wavelengths set at 460nm and 483nm respectively.

### Mass Spectrometry

The fractions of the cellular samples from the GELFrEE^®^ system were passed through HiPPR™ Detergent Removal 0.1ml columns (ThermoFisher Scientific), previously equilibrated with a 25mM ammonium bicarbonate (NH_4_HCO_3_) solution, to remove the detergent from the solution. The samples were centrifuged for 2mins at 1500g, then lyophilized.

Samples were then resuspended in 50mM ammonium bicarbonate before being first reduced (10mM dithiothreitol) for 40mins at 56 °C, alkylated (20mM iodoacetamide) for 30mins at room temperature, and finally digested with trypsin for 16hours at 37 °C (1/50 w/w enzyme/proteins ratio). Reactions were stopped by acidifying the solution using 10% TFA. Generated peptides were then analyzed by LC-MS/MS.

Peptides were separated by reversed-phase chromatography using Ultra Performance Liquid Chromatography (UPLC-MClass, HSS T3 column, Waters, Milford, MA, USA) in one dimension with a linear gradient of acetonitrile (5 to 40% in 70mins, solvent A was water 0.1% formic acid, solvent B was acetonitrile 0.1% formic acid) at a 600 nl/min flow rate. The chromatography system was coupled with a Thermo Scientific Q Exactive™ Plus hybrid quadrupole-Orbitrap™ mass spectrometer (ThermoFisher Scientific, Waltham, MA, USA), with a targeted method. The targeted masses are the m/z of the three following doubly charged peptides, that result from the trypsin digestion: LVFFAEDVGSNK m/z 663.3404, GAIIGLMVGGVV m/z 543.3230 and GAIIGLMVGGVVIA m/z 635.3836.

Full-MS scans were acquired at 70.000 mass resolving power (full width at half maximum). A mass range from 400 to 1750 m/z was acquired in MS mode, and 3×10^6^ ions were accumulated. Ion trap Higher energy Collision Dissociation fragmentations at NCE (Normalized Collision Energy) 25 were performed within 2amu isolation windows.

Raw MS files were analyzed by Proteome Discoverer 2.1.1.21 software (Thermo Scientific). MS/MS spectra were compared to the Uniprot *Cricetulus griseus* protein database, in which had been added the sequence of two main human amyloid peptides Aβ40 and Aβ42. Due to this restricted length in amino acids, the criteria for identification of each protein has been set up to one unique peptide per protein. The false discovery rate (FDR) was set to 0.01 on both protein and peptide levels. The tolerance on mass accuracy was set at 5 ppm (10 ppm for MS/MS).

### qPCR

RNAs were extracted from cells in TRIPure^®^ reagent and reverse-transcribed using an iScript cDNA synthesis kit. qPCR conditions were 95°C for 30secs, followed by 40 cycles of 30secs at 95°C, 45secs at 60°C and 15secs at 79°C and ended by 1 cycle of 15secs at 79°C and 30secs at 60°C. The relative changes in the target gene-to-GAPDH mRNA ratio were determined by the 2^(−ΔΔCt)^ calculation. The sequences for qPCR primers are provided in Supplemental Table S3.

### Data Availability

The datasets generated during and/or analysed during the current study are available from the corresponding author on reasonable request.

## Acknowledgments

The authors thank Esther Paître and Pierre Burguet for their technical support. We also thank Loic Quinton for kindly providing us with the Gel Eluted Liquid Fraction Entrapment Electrophoresis (GELFrEE) 8100 system. Funding to P.K.C is acknowledged from SAO-FRA Alzheimer Research Foundation, Fondation Louvain and Queen Elisabeth Medical Research Foundation (FMRE to P.K.C and LQ). The work was supported by funds from FNRS grant PDRT.0177.18 to P.K.C and LQ.

## Author Contributions

D.M.V designed and performed research, analysed data and wrote the manuscript. C.V performed research and analysed data. P.B performed mass spectrometry experiments under the supervision of L.Q. N.C and S.C analysed and interpreted data. L.C.S kindly provided us with the vAβ sequence and provided guidance with interpretation of data. P.K.C. supervised, designed research project and interpretations. All authors reviewed the manuscript.

## Additional Information

The authors declare no competing interests. Correspondence and requests for materials should be addressed to P.K.C.

## Notes

### Competing Interest Statement

The authors have declared no competing interest.

