## Supplemental Data for "Cell-derived hexameric β-amyloid: a novel insight into composition, self-assembly and nucleating properties"

[†]Current affiliation

***Corresponding author:**

**Pascal Kienlen-Campard**

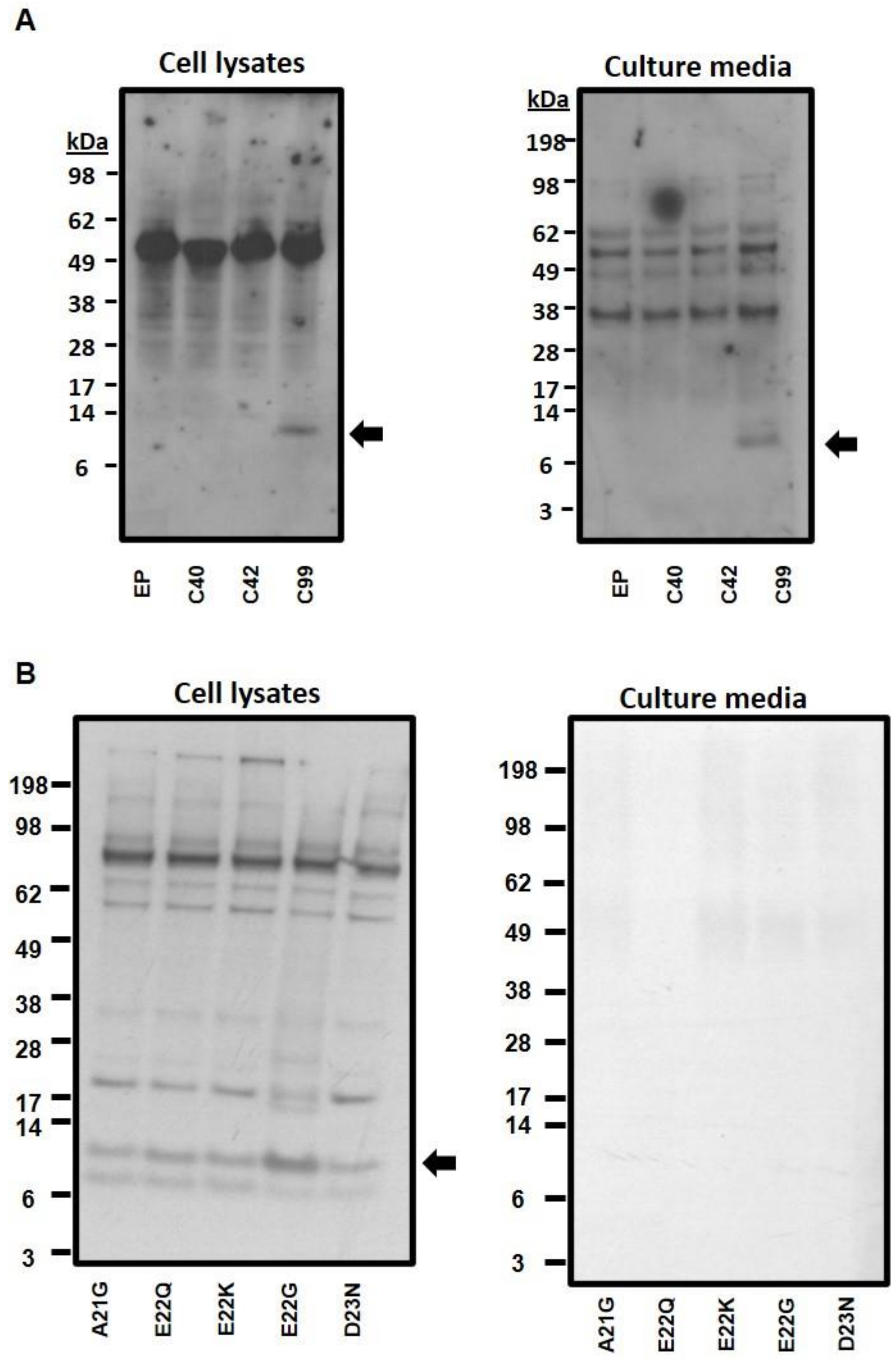


**Supplemental Figure 1. APP C-terminal detection with C-ter antibody confirms assemblies to be Aβ. (A)** Hexameric assemblies are not detected in the cell lysates or culture media of CHO cells transfected with EP, C40, C42 or C99 **(B)** FAD mutants. The C-terminal fragment is seen in the cell lysates of CHO cell transfected with C99 sequences (black arrows).

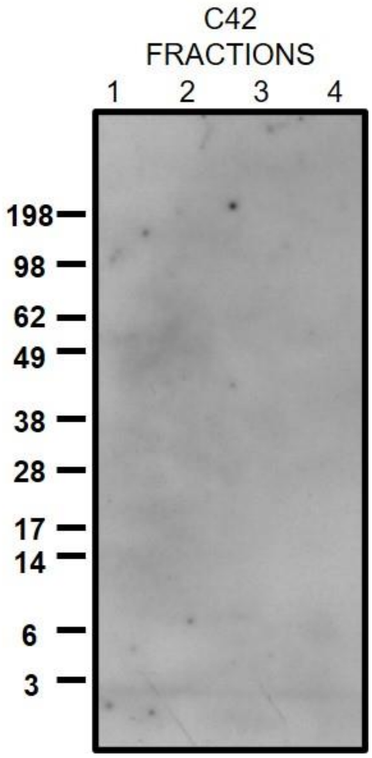


**Supplemental Figure 2. Isolation of C42 derived hexamers: Fractions 1-4.** No Aβ assemblies were isolated in fraction 1-4 collected after separation by the GELFrEE 8100 system. Western blot detected with the W02 antibody.

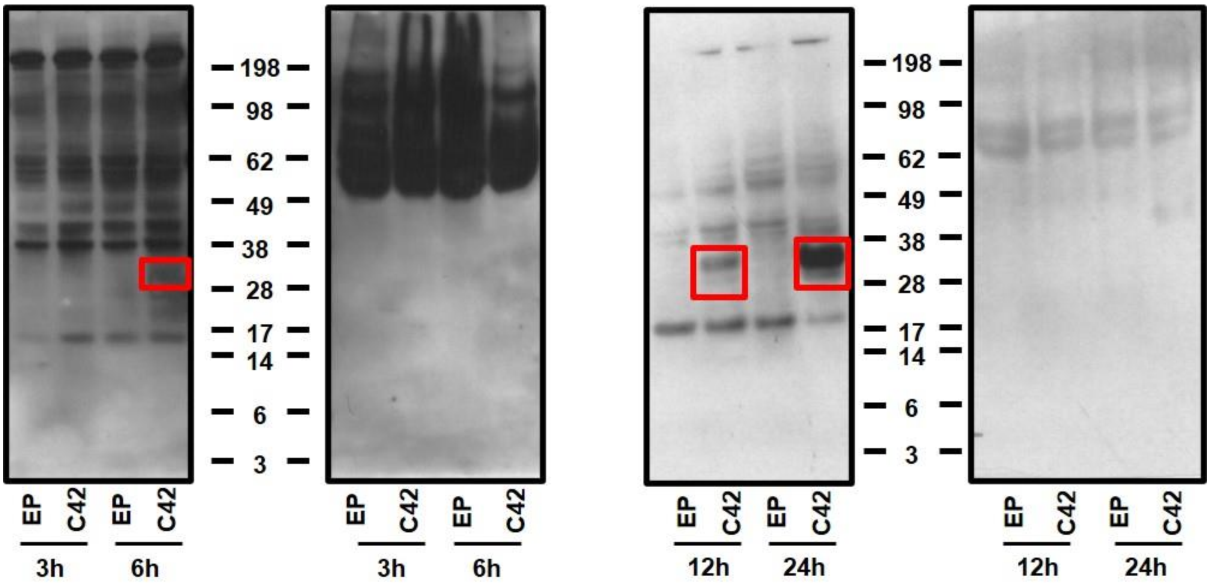


**Supplemental Figure 3. Lower molecular weight Aβ assemblies were not detected in C42 transfected CHO cells.** CHO cells were transfected with C42 for 3,6,12 and 24hours. Formation of hexameric assemblies were seen in cell lysates by 6 hours with no lower weight assemblies seen at the earlier time point of 3 hours. Aβ assemblies were not detected in the media of these cells even after 24 hours. Western blot detection with the W02 antibody.

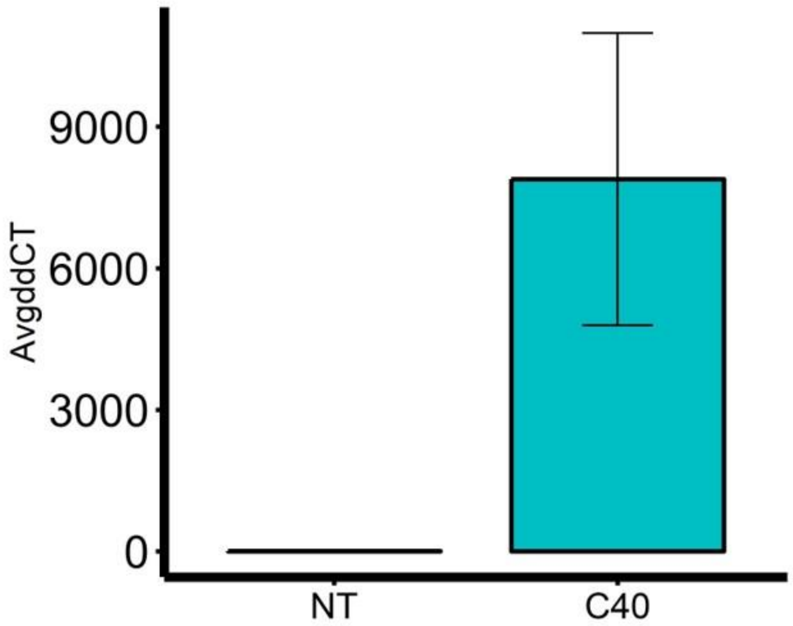


**Supplemental Figure 4. C40 transfection efficiency.** qPCR confirms that the lack of hexameric assemblies is not due to low transfection efficiency (N=5)

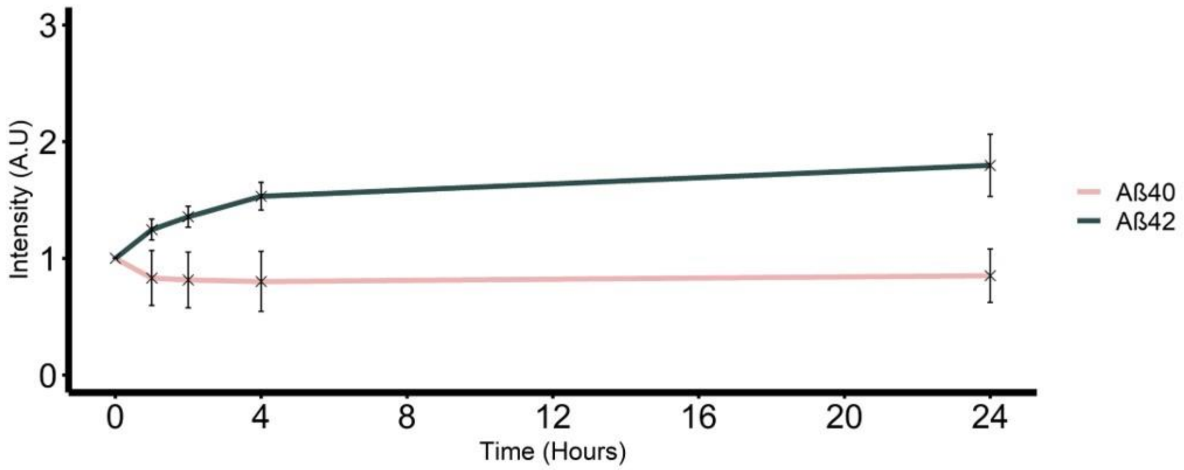


**Supplemental Figure 5. ThT fluorescence of Aβ40 and Aβ42 over 24 hours.** 50µM of each peptide was incubated with 20µM ThT and fluorescence intensity was measured over 24hours. Aβ40 does not display an increase in fluorescence however Aβ42 displays a sharp increase in fluorescence intensity from 0-4hours which then plateaus over 24 hours.

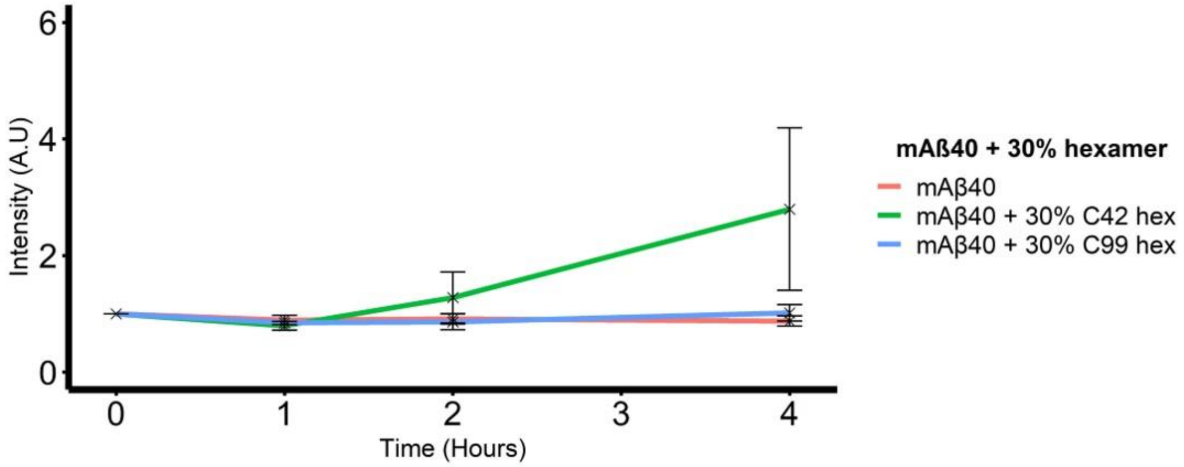


**Supplemental Figure 6. monomeric Aβ40 does show accelerated aggregation after 30% hexamer addition.** 30% addition both hexamers leads to increased ThT fluorescence over 24 hours.

| | **Sequence** |
|---|---|
| C42 | DAEFRHDSGYEVHHQKLVFFAEDVGSNKGAIIGLMVGGVVIA |
| vC42 | DAEFRHDSGYEVHHQKLV**S**FAEDVGSNKGAIIGLMV**D**GVVIA |

Supplemental Table S1. **Primary sequence of the wild type (C42) and variant C42 (cvC42) sequences**

| | Forward | Reverse |
|---|---|---|
| A21G | 5'-ggtgttctttggagaagatgtgggttcaaac-3' | 5'-cacatcttctccaaagaacaccaatttttgatg-3' |
| E22Q | 5'-ggtgttctttgcacaagatgtgggttcaaac-3' | 5'-cccacatcttgtgcaaagaacaccaatttttg-3' |
| E22K | 5'-ggtgttctttgcaaaagatgtgggttcaaac-3' | 5'-cccacatcttttgcaaagaacaccaatttttg-3' |
| E22G | 5'-gtgttctttgcaggagatgtgggttcaaac-3' | 5'-cccacatctcctgcaaagaacaccaatttttg-3' |
| D23N | 5'-ggtgttctttgcagaaaatgtgggttcaaac-3' | 5'-cccacattttctgcaaagaacaccaatttttg-3' |

Supplemental Table S2. **Primer sequences for site directed mutagenesis of C99 to introduce FAD mutations**

| | Forward | Reverse |
|---|---|---|
| GAPDH | 5’-acccagaagactgtggatgg-3’ | 5’acacattgggggtaggaaca-3’ |
| C40 | 5’ gctggaggatgcagaattccgac 3’ | 5’ cgcccaccatgagtccaatgattg 3’ |

Supplemental Table S3. **Primer sequences for qPCR to probe for C40 transfection efficiency in CHO cells.**
